# Oxytocin Receptor Expression and Activation in Parasympathetic Brainstem Cardiac Vagal Neurons

**DOI:** 10.1101/2025.03.19.644204

**Authors:** Xin Wang, Caitlin Ribeiro, Anna Nilsson, Joan B. Escobar, Bridget R. Alber, John R. Bethea, Vsevolod Y Polotsky, Matthew W. Kay, Kathryn Schunke, David Mendelowitz

## Abstract

Autonomic imbalance, particularly reduced activity from brainstem parasympathetic cardiac vagal neurons (CVNs) is a major characteristic of many cardiorespiratory diseases. Therapeutic approaches to selectively increase CVN activity have been limited by lack of identified selective translational targets. Recent work has shown that there is an important excitatory synaptic pathway from oxytocin (OXT) neurons in the paraventricular nucleus of the hypothalamus (PVN) to brainstem CVNs, and that OXT could provide a key selective excitation of CVNs. In clinical studies, intranasal OXT increases parasympathetic cardiac activity, autonomic balance, and reduces obstructive event durations and oxygen desaturations in obstructive sleep apnea patients. However, the mechanisms by which activation of hypothalamic OXT neurons, or intranasal OXT, increases brainstem parasympathetic cardiac activity is poorly understood. CVNs are located in two cholinergic brainstem nuclei: the nucleus ambiguus (NA) and dorsal motor nucleus of the vagus (DMNX). In this study we characterize the co-localization of OXT receptors in CVNs (OXTR), as well as non-CVN cholinergic neurons, located in the NA and DMNX nuclei. Selective chemogenetic excitation of OXTR+ CVNs was performed by expressing DREADDs with a combination of Cre and flp dependent viruses. We found that OXT receptors are highly expressed in CVNs in the DMNX and OXT increases DMNX CVN activity, but the receptors and responses are absent in CVNs in the NA. Selective chemogenetic activation of OXTR+ CVNs in the DMNX evoked a rapid and sustained bradycardia.

## 1 INTRODUCTION

Autonomic imbalance, particularly reduced activity from brainstem parasympathetic cardiac vagal neurons (CVNs) is a major characteristic of many cardiorespiratory diseases including hypertension, sudden cardiac death, sleep apnea, heart failure, sudden unexpected death in epilepsy as well as other diseases such as obesity, anxiety and osteoarthritis (Sohn et al. (2024), Kay et al. (2022), Jones et al. (1998), Strain et al. (2023), Cheshire (2013), Strain et al. (2024)). Therapeutic approaches to selectively increase parasympathetic cardiac activity have been limited by lack of identified translational targets in CVNs. Recent work has shown that there is an important excitatory synaptic pathway from oxytocin (OXT) neurons in the paraventricular nucleus of the hypothalamus (PVN) to brainstem CVNs, and that this pathway provides a key excitation to CVNs that are inherently silent (Pinol et al. (2012), Mendelowitz (1996)). Selective chemogenetic activation of PVN OXT neurons, using Designer Receptors Exclusively Activated by Designer Drugs (DREADDs), reduces heart rate (HR) and blood pressure by increasing cardiac vagal tone (Dyavanapalli et al. (2020b)). Activation of PVN OXT neurons, as well as intranasal OXT, increases parasympathetic activity to the heart, restores autonomic balance, and prevents or mitigates the adverse cardiorespiratory dysregulation that occurs in many diseases with autonomic imbalance, including chronic intermittent hypoxia induced hypertension (Rodriguez et al. (2023)), heart failure (Garrott et al. (2017), Dyavanapalli et al. (2020a), Dyavanapalli et al. (2020b)), and myocardial infarction (Schunke et al. (2023)). In clinical studies, intranasal OXT increases parasympathetic cardiac activity, autonomic balance, and reduces obstructive event durations and oxygen desaturations in obstructive sleep apnea patients (Jain et al. (2017), Jain et al. (2020)). However, the mechanisms by which activation of hypothalamic OXT neurons, or intranasal OXT, increases brainstem parasympathetic cardiac activity is poorly understood. CVNs are located in two cholinergic brainstem nuclei: the nucleus ambiguus (NA) and dorsal motor nucleus of the vagus (DMNX). In this study we characterize the co-localization of OXT receptors in CVNs (OXTR), as well as non-CVN cholinergic neurons, located in the NA and DMNX nuclei. Selective chemogenetic excitation of OXTR+ CVNs was performed by expressing DREADDs with a combination of Cre and flp dependent viruses. We found that OXT receptors are highly expressed in CVNs in the DMNX and OXT increases DMNX CVN activity, but the receptors and responses to OXT are absent in CVNs in the NA. Selective chemogenetic activation of OXTR+ CVNs in the DMNX evoked a rapid and sustained bradycardia.

## 2 METHODS

### 2.1 Animals

All animal experiments were performed in accordance with NIH and Institutional Animal Care and Use Guidelines, approved by the George Washington University Institutional Animal Care and Use Committee (IACUC, protocol #2022-028). The transgenic OXT receptor (OXTR)-Cre mouse line (B6.Cg-*Oxtr*^*tm*1.1(*cre*)*Hze*^/J, Common Name: Oxtr-T2A-Cre-D, Jackson Labs stock # 031303) and the Cre dependent floxed ChR2-eGFP mouse line (ROSA26-EGFP Cre recombinase report mouse, Jackson Labs stock # 012569) were used in the study. Mice were housed at standard environmental conditions (24°C–26°C in the 12-h light/dark cycle, 7a.m.–7p.m. lights on), until the experiments started. Water and food were available *ad libitum*.

Mice were crossbred to express ChR2-eYFP in OXTR+ neurons. One parent (OXTR-Cre mouse) had homozygous expression of an OXTR promoter to guide the expression of Cre recombinase while the other parent (Cre dependent floxed ChR2-eYFP mouse) had homozygous ChR2-eYFP fusion protein expression dependent upon Cre expression. Offspring (OXTR-ChR2-EYFP) were genotyped by Transnetyx before experiments.

### 2.2 Retrograde labeling of preganglionic parasympathetic cardiac vagal neurons (CVNs)

To label CVNs in the NA and DMNX, cholera toxin B (CTB) conjugated to Alexa Fluor 555 (Invitrogen C22843) was injected into the right atrial pericardial space of postnatal day 5 (P5) OXTR-ChR2-EYFP mice. Following injection, mice recovered on a 37°C heating pad until regaining the righting reflex, after which they were returned to the litter. 3-7 days post-injection, CVNs in the brainstem NA and DMNX nuclei were identified by presence of fluorescent tracer (red; CTB-AlexaFluor™ 555).

### 2.3 Selective DREADDs expression in OXTR+ CVNs

To selectively express DREADDs in DMNX OXTR+ CVNs we used a two-stage combination of Cre and FlpO expressing viruses in a separate group of mice (n=7). A retrograde Cre dependent site-specific recombinase FlpO virus (AAVrg pEF1a-DIO-FLPo-WPRE-hGHpA, addgene # 87306; 1 µL, ≥ 7×10^12^ vg/mL) was injected into the pericardial sac of P5 OXTR-ChR2-EYFP mice (same procedure as above) in an initial surgery. Four weeks later, the Flp-dependent excitatory hM3D (Gq) DREADDs mCherry virus (pAAV-hSyn-fDIO-hM3D(Gq)-mCherry-WPREpA (AAV8), ≥ 1×10^13^ vg/mL, Addgene viral prep # 154868-AAV8) was injected into the DMNX. Briefly, mice were anesthetized using a mixture of ketamine (100 mg/kg) and xylazine (10 mg/kg), then secured in a stereotaxic frame (David Kopf Instruments, CA, USA). Using stereotaxic guidance of a pulled glass capillary (40µm tip diameter) (G1, Narishige, UK) attached to a pneumatic microinjector (IM-11-2, Narishige, UK), 240 nL of viral preparation was bilaterally injected into the DMNX (AP 0, at the level of Obex, ML 0.15mm, DV 0.5mm). The capillary remained in place for 5 minutes to allow diffusion, and was then removed slowly to avoid dispersion to neighboring brainstem regions. DREADDs expressing DMNX parasympathetic CVNs were identified by the presence of mCherry.

### 2.4 Brain slice preparation, Immunohistochemistry, and RNAScope in situ hybridization

One week after injecting Alexa Fluor 555-conjugated CTB into the pericardial sac, P12–P14 pups were anesthetized by isoflurane and transcardially perfused with phosphate buffered saline (PBS) followed by 4% paraformaldehyde (PFA). Brains were dissected, post-fixed in 4% PFA overnight at room temperature, then washed 3×10 min in PBS. A serial of 50 µm coronal section medullary slices containing the NA and/or DMNX were obtained using a Leica dissection vibratome (Leica VT 1000S). 10-20 NA and 9-11 DMNX medullary slices were obtained from each animal. All slice sections were processed using immunohistochemistry for choline acetyltransferase (ChAT) and eYFP to identify cholinergic and OXTR-expressing neurons, respectively. After blocking nonspecific proteins in 10% normal goat serum (NGS) in PBS with 0.3% Triton X-100 (PBST) for 4 hours at room temperature, slices were incubated in primary antibody at 4°C for 24 hours (mouse monoclonal-ChAT, diluted 1:500, ThermoFisher, Cat# MA5-31382, Waltham, MA; and chicken anti-GFP/EYFP, 1:1500 dilution; Abcam, ab 13970, Cambridge, MA). Secondary antibodies were applied for 4 hours at room temperature. Secondary antibodies were goat anti-mouse antibody conjugated with Alexa Fluor 647 (ThermoFisher Cat # A-21241) and goat anti chicken Alexa Fluor 488 (ThermoFisher Cat A 11039), both 1:200 dilution. Slides were mounted with Fluorogel (Electron Microscopy Sciences, SKU: 17985-10) and imaged with a Leica TCS SP8 multi-photon confocal microscope equipped with supercontinuum white laser source and single molecule detection hybrid detectors (SMD HyD, Leica, Wetzlar, Germany).

For RNAscope in situ hybridization (ISH) experiments, animals were anesthetized by isoflurane and transcardially perfused with RNAse-free PBS, followed by 4% PFA. Brain tissues were obtained and fixed in 4% PFA solution for 4 hours at 4°C. After fixation, brains were transferred into RNAse free 30% sucrose (PBS, for dehydration and cryoprotection) and stored at 4°C for 48 hours, then stored at −80°C and cryosectioned coronally 12µm thick. Every 4^th^ section was prepared according to the manufacturer’s instructions, using the RNAscope Multiplex Fluorescent Detection Reagents v2 kit (ACDBio, 323110) combined with Integrated Co-Detection (ACDBio, 323180). Briefly, slides were baked at 60°C, fixed in 4% PFA, and dehydrated. Slides were incubated in Hydrogen Peroxide (ACDBio, 322335), followed by antigen retrieval using RNAscope 1X Co-Detection Target Retrieval buffer (ACDBio 323165) heated to 99°C. Primary antibody to detect OXTR containing neurons (Chicken anti-GFP/EYPF; Abcam, AB13970, 1:1500) was incubated overnight at 4°C. Slides were post-fixed using 10% NMBF followed by the RNAscope Multiplex Fluorescent Detection protocol according to the manufacturer’s instructions. RNAscope Probe C1 mmOxtr (ACDBio, 412171) was used to target OXT receptor mRNA and Vivid Fluorophore 650 (ACDBio, 323273, 1:500) was used to label the OXTR probe. Finally, slides were incubated in secondary antibody AlexaFluor Goat anti-chicken 488 (Invitrogen, A11039, 1:200), stained with DAPI (ACDBio, 323108), mounted with Prolong Gold Antifade Mountant (Invitrogen, P36934), and stored at 4°C until imaging.

### 2.5 Patch clamp electrophysiology experiments

To study the effect of OXT on CVN activity, OXTR-ChR2-EGFP mice received CTB pericardial sac injections at P5, and whole-cell patch clamp recordings of CVNs occurred at P8-P12. Animals were anesthetized with isoflurane, sacrificed, and transcardially perfused with ice-cold NMDG aCSF (in mM): 93 NMDG, 93 HCl, KCl, 1.2 NaH_2_PO_4_, 30 NaHCO_3_, 25 glucose, 20 HEPES, 5 Sodium Ascorbate, 2 Thiourea, 3 Sodium pyruvate, 10 MgSO_4_.7H_2_O, 0.5 CaCl_2_.2H_2_O, oxygenated with 95% O_2_ and 5% CO_2_, (pH 7.4). The brain was carefully removed and brainstem slices (250 µm) were prepared using a vibratome. Tissue slices were then moved from the bath solution and incubated in the NMDG recovery solution (in mM): 93 NMDG, 93 HCl, 2.5 KCl, 1.2 NaH_2_PO_4_, 25 NaHCO_3_, 20 HEPES, 25 D-Glucose, 10 MgSO_4_, 0.5 CaCl_2_ bubbled with 95% O_2_/5% CO_2_ at 34°C for 10 min. Tissue slices were then moved to a recording chamber and perfused with standard aCSF solution containing (in mM): 125 NaCl, 3 KCl, 25 NaHCO_3_, 5 HEPES, 5 D-glucose, 1 MgSO_4_, 2 CaCl_2_ and continuously bubbled with 95% O_2_/5% CO_2_ to maintain pH at 7.4 at room temperature (22°C–24°C). Patch electrodes were filled with an intracellular recording solution at pH 7.3 containing (in mM): 130 K-Gluconate, 4 KCl, 10 HEPES, 0.3 EGTA, 4 Mg-ATP, 0.3 Na_2_-GTP, 10 phosphocreatine-Na_2_, and 13.4 Biocytin. Biocytin (0.5%, a highly photostable far-red (excitation laser 640nm) biocytin fluorophore conjugated dye, CF640R, Biotium, Inc., Fremont, CA, USA) was added in the patch solution to further identify the neurons in the NA and DMNX using immunohistochemistry staining. Identified CVNs were voltage-clamped at a holding potential of −80 mV. Gabazine (25 µM) was used to block GABAergic inhibitory neurotransmission, and strychnine (1 µM) was included to block glycinergic inhibitory neurotransmission. Focal drug application was performed using a PV830 Pneumatic PicoPump pressure delivery system (WPI, Sarasota, FL). Drugs were ejected from a patch pipette positioned within 30 µm from the patched CVN. The maximum range of drug application has been previously determined to 100-120 µm downstream from the drug pipette, and considerably less behind the drug pipette (Wang et al. (2001)). Electrophysiological data were digitized and collected via Clampex (10.2) and analyzed using Clampfit (10.7).

### 2.6 Confocal Imaging

Leica TCS SP8 MP confocal microscopy was used to assess colocalization of ChR2-eGFP labeled OXTR+ neurons, CVNs, and ChAT (cholinergic) neurons in the DMNX and NA. Tissue slices containing the DMNX and NA were examined with 418, 514 and 610 nm wavelengths to visualize eGFP, CTB-Alexa Fluor 555, and ChAT-Alexa Fluor 647, respectively. Images were captured with a DFC365FX camera at 2048 by 2048-pixel resolution. Full field images of the entire slice were taken at 10x to localize the NA and DMNX. Z-stacks were then taken with 20x/0.75 oil-immersion objective, at z-step size of 0.9 µm to produce image volumes allowing for the identification of colocalization of ChAT, CVNs and ChR2-eGFP positive neurons. Images were processed and analyzed using Imaris 10 software (Oxford Instruments Oxford, UK). Brain slices for RNAscope ISH were imaged with the automated Leica MDI8 Thunder microscope mounted with a Hamamatsu ORCA-Flash 4.0 v3 camera and paired with advanced LASX software for OXTR mRNA detection. A 63X/1.40 oil immersion objective was used to image processed sections, illuminated with the LED8 spectrum to visualize GFP-eYFP-488, OXTR mRNA-Vivid 650, CTB-Alexa Fluor 555, and DAPI. A total of 6-10 sections spanning the DMNX from 2 mice were analyzed. Rostral sections of the NA in the brainstem showed no OXTR positive cells and were not quantified further.

### 2.7 In vivo Heart Rate Measurements

One month after DREADDs injection, animals were implanted with a telemetry device (DSI wireless transmitters, ETA-F10 [Data Sciences International, St. Paul, Minnesota]) with electrocardiographic (ECG) leads to measure HR. One-week later, animals were acclimated to the whole-body plethysmography system (Scireq, Montreal, QC Canada) for 3 hours on 3 consecutive days. Control ECG and respiratory rate (RR) were recorded following an IP injection of either saline or Clozapine-n-Oxide (CNO) (1 mg/kg) to activate excitatory DREADDs in OXTR+ CVNs. HR data were obtained by analyzing the raw ECG signal using the ECG analysis platform of Labchart (ADInstruments, Colorado Springs, CO). RR data were obtained using EMKA IOX Software and analyzed using the same approaches.

### 2.8 Data Analysis

All data are presented as mean with 95% CI. Statistical comparisons were made using repeated-measures (RM) one-way and two-way ANOVA with Dunnett’s post-tests, as well as paired or unpaired Student’s t tests, as appropriate. A p-value *<* 0.05 was considered statistically significant. Statistical analyses were conducted using GraphPad Prism (Graphpad Software, San Diego, CA).

## 3 RESULTS

CVNs were nearly equally distributed in the NA and DMNX nuclei (467 *±* 97 CVNs in the DMNX and 439 *±* 53 CVNs in the NA per animal). As expected, CVNs in both nuclei were highly co-localized with ChAT (93 *±* 1% in the DMNX and 95 *±* 1% in the NA, n=7 and n=8, respectively). CVNs comprised approximately 40% of the ChAT population in these nuclei, with the population of CVNs consisting of 38.5 *±* 3.8% of the ChAT neurons in the DMNX, and 44.8 *±* 2.9% of the ChAT neurons in the NA, figure 1.

**Figure 1.**
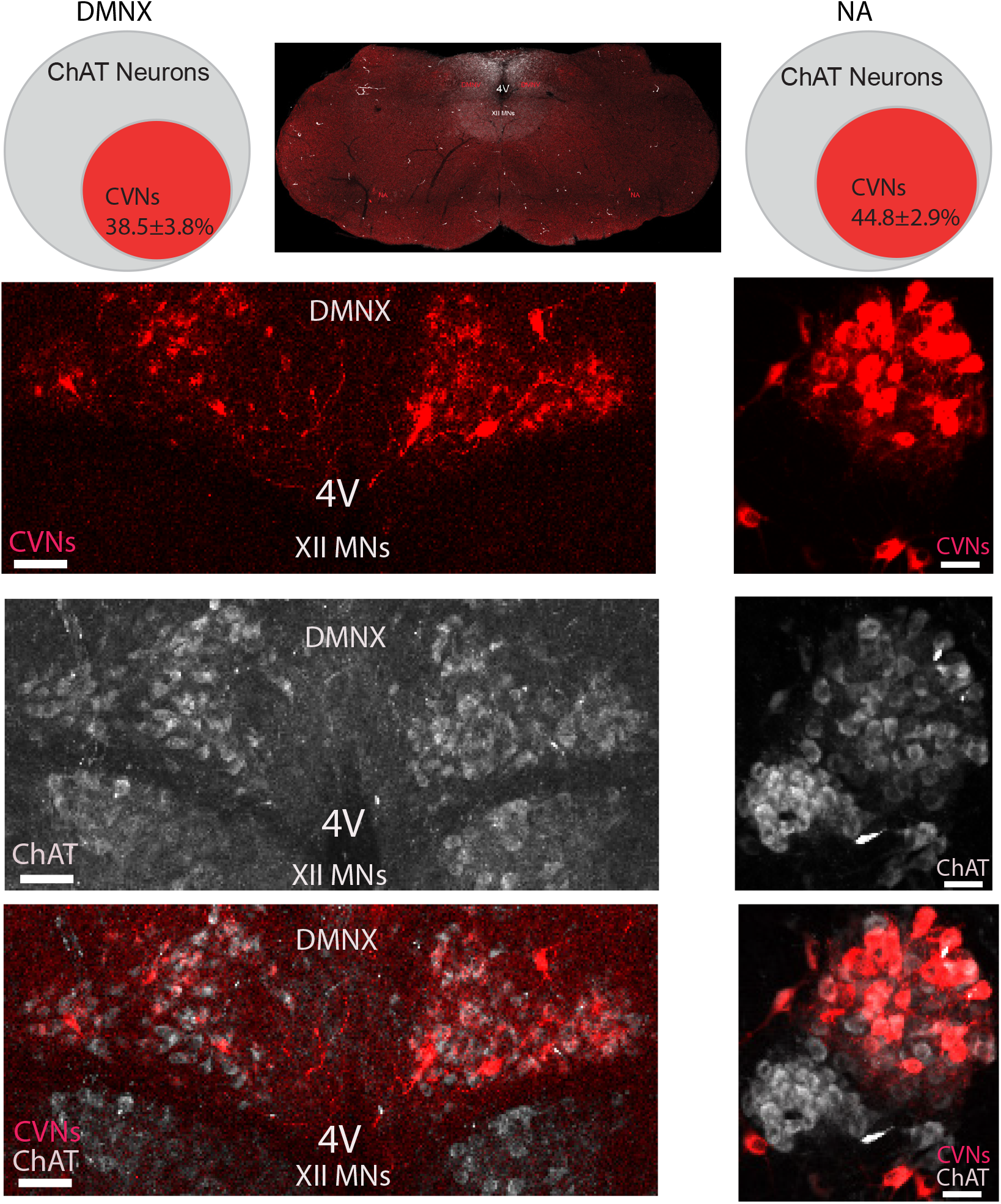
Percentage of CVNs-ChAT neurons in the DMNX and NA in OXTR-Cre-ChR2-eYFP mice. Top middle: An example confocal image showing overview of brainstem containing CVNs (red) in the DMNX and NA. The DMNX data presented on the left and NA on the right panel. The top two Venn diagrams showing CVNs account for the percentage population of the cholinergic neurons in the DMNX (left) and NA (right). Example confocal images showing the CVNs in red (top), ChAT in grey (middle), and merged CVNs and ChAT neurons light red (bottom) in the DMNX and NA, respectively. Scale bar: 50 µm. For this and all the following figures: 4V, the 4^th^ ventricle; XII MNs, hypoglossal motorneurons; CVNs, cardiac vagal neurons; ChAT, cholinergic neurons.

OXTR+ neurons were highly prevalent within the DMNX (746 *±* 199 OXTR+ neurons/animal) but were sparse in the NA (74 *±* 14 OXTR+ neurons/animal, n=8), as illustrated in figure 2. A majority of CVNs in the DMNX were OXTR+ as 54 *±* 3% of CVNs in the DMNX were also OXTR+, see figure 2. Some non-CVN DMNX neurons also co-localized with OXTR+, 24.8 *±* 4.9%. In contrast, very few CVNs in the NA were OXTR+ neurons (1.5 *±* 0.2%) and similarly a small number of non-CVN ChAT neurons in the NA were OXTR+ neurons (1.5 *±* 0.2%), figure 3.

**Figure 2.**
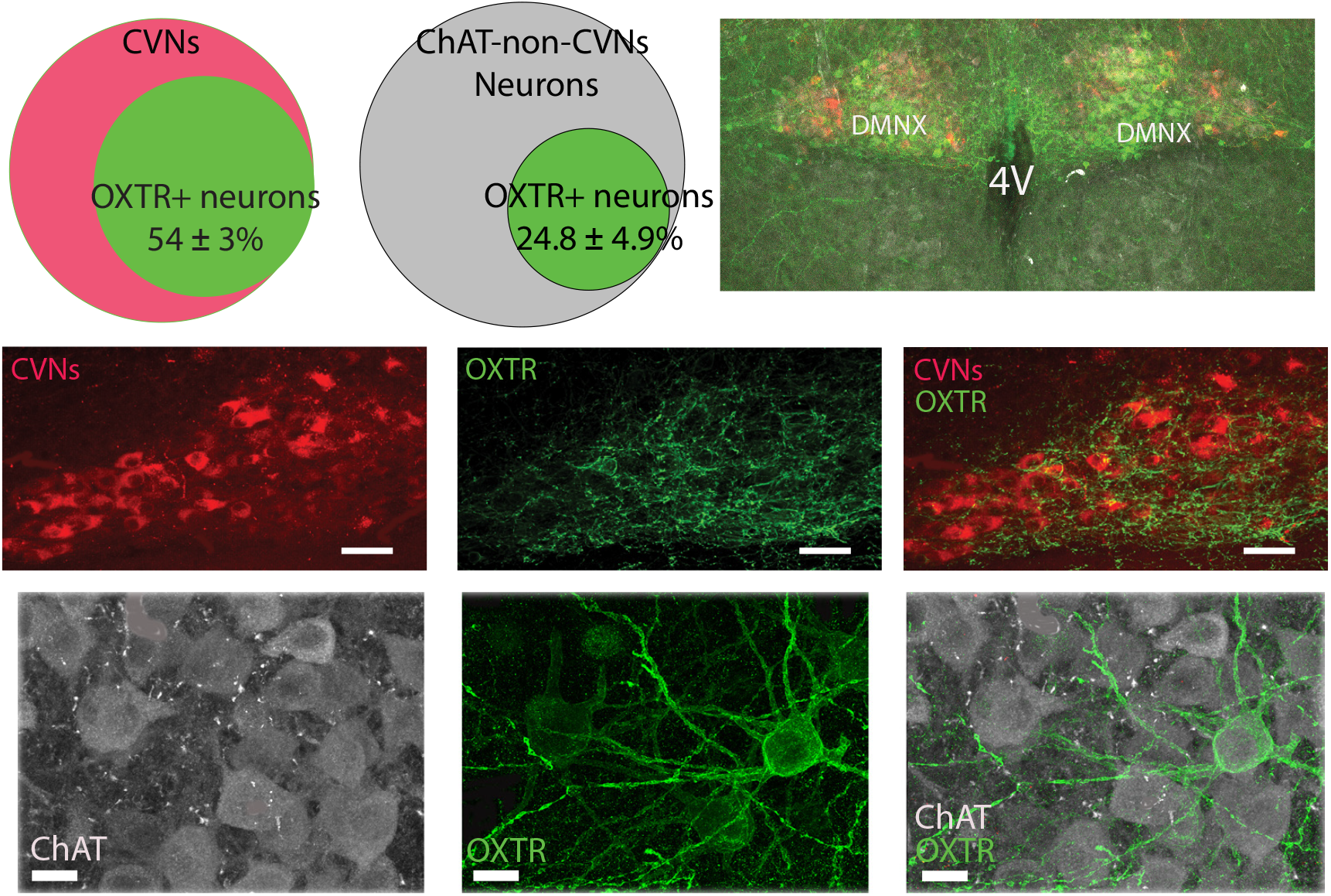
Colocalization of OXTR+ neurons with CVNs and non-CVN ChAT cells in the DMNX. Top right: an example of a low-resolution confocal image showing overview of brainstem dorsal side DMNX area containing CVNs (red), ChAT (grey), and OXTR+ (green). The middle panel represents CVNs colocalization with OXTR+ neurons: Venn diagram showing the percentage of CVNs containing OXTR; the confocal images mark CVNs in red (left), OXTR+ in green (middle), and CVNs merged with OXTR+ labeling (right); scale bar 20 µm. The bottom panel illustrates the OXTR+ neurons colocalized with ChAT neurons that are not CVNs. The Venn diagram showing the percentage of non-CVN ChAT neurons containing OXTR; Typical confocal images illustrate ChAT neurons in grey (left), OXTR+ neurons in green (middle), and merged OXTR+ and ChAT neurons (right); scale bar: 10 µm. OXTR: Oxytocin receptor

**Figure 3.**
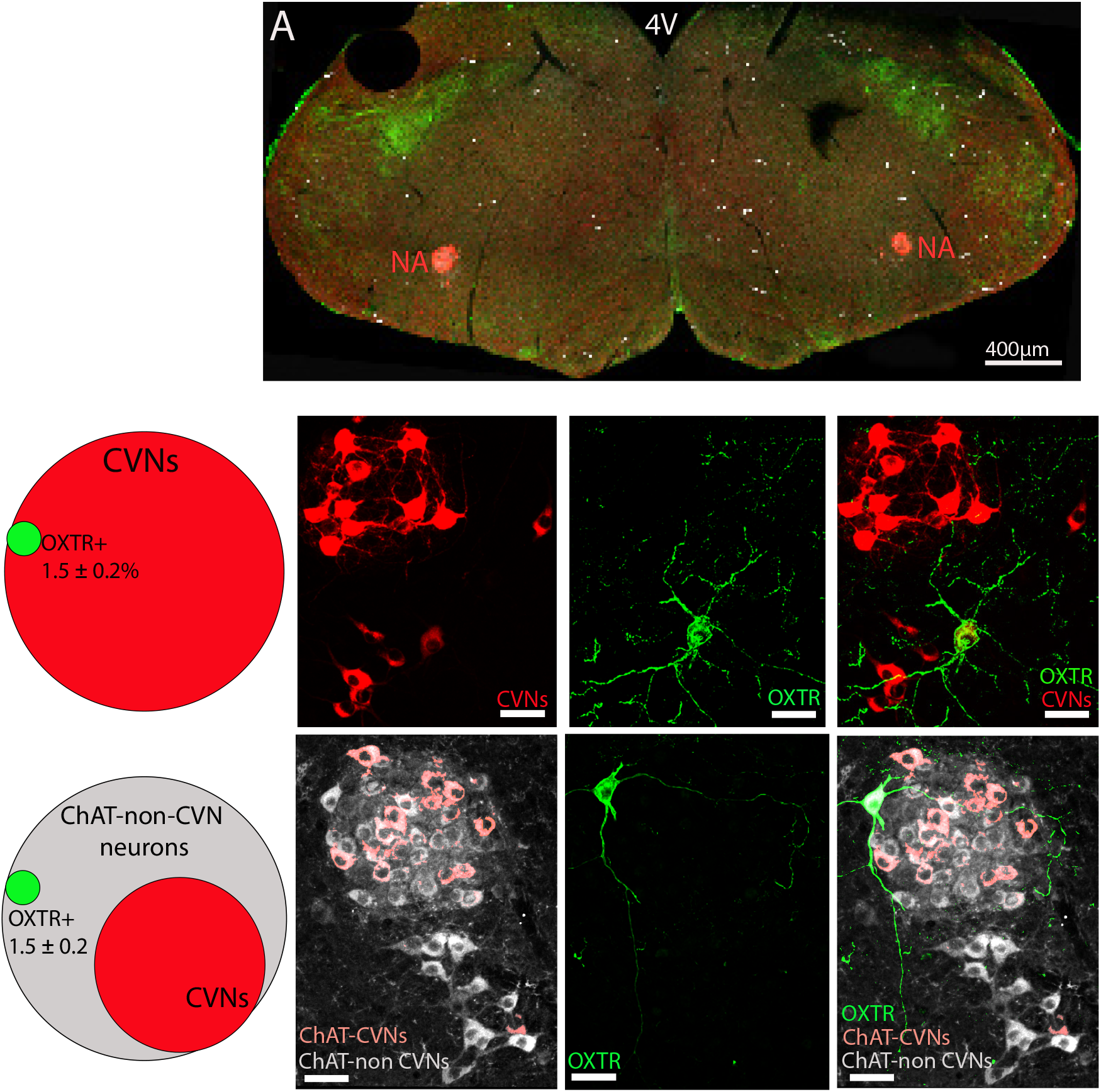
The OXTR+ neurons colocalization with CVNs and non-CVN ChAT cells in the NA. A: The low-resolution confocal image showing overview of a brainstem containing CVNs (red), ChAT (grey), and OXTR+ (green) in the NA. The middle panel reveals the percentage of CVNs containing OXTR+ (Venn diagram on the left), confocal images represent CVNs in red (2^nd^ left), OXTR+ neurons in green (2^nd^ from right), and colocalized OXTR+ neuron with CVNs (right). The bottom panel denotes the percentage of non-CVN ChAT neurons containing OXTR+ (Venn diagram on the left), confocal images showing ChAT in grey (2^nd^ left), OXTR+ neurons in green (2^nd^ from right), and colocalized OXTR+ neuron with non-CVN ChAT neurons (right). Scale bar: 50 µm.

## 3.1 ChR2-eGFP expressing neurons are correlated with active OXTR mRNA transcription

To test whether the crosssbred OXTR+cre x floxed ChR2-eGFP neurons express OXTR mRNA as adults, we performed RNAscope ISH experiments in three adult OXTR-Cre-ChR2-eGFP mice. As shown in figure 4, eGFP labeled OXTR+-cre ChR2 positive neurons (green) were highly co-localized with the OXTR mRNA probe (red) with 98.6% of OXTR mRNA+ cells also expressing ChR2-eGFP+. Additionally, 100% of the OXTR-Cre-ChR2-eGFP positive neurons were co-localized with OXTR mRNA. No OXTR mRNA+ neurons were found in the NA.

**Figure 4.**
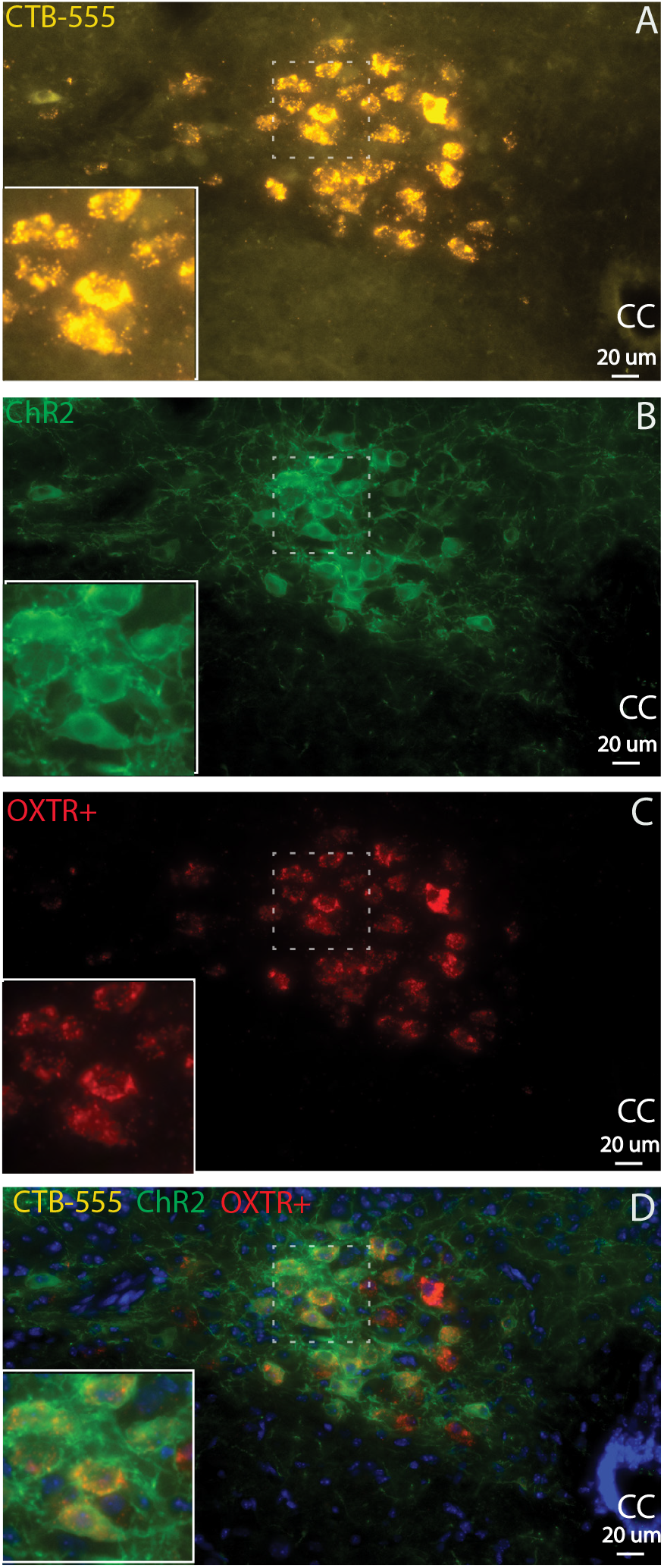
ChR2-eGFP expressing neurons are correlated with active OXTR mRNA transcription. A: CTB-555 labeled CVNs (gold). B: ChR2-eGFP labeled OXTR^+^ neurons and fibers (green). C: OXTR mRNA+ cells (red). D: All of OXTR^+^-Cre ChR2 positive neurons (green) co-localized with the OXTR mRNA probe (red). Insert: focus on the dash line marked area. cc, central canal.

## 3.2 Changes in firing in OXTR+ CVNs with OXT

To test the effect of OXT on OXTR+ CVNs activity, patch clamp recordings were obtained to quantify firing rates before and after OXT administration. Application of OXT (100 nM) significantly increased the firing rate of CVNs in the DMNX by 20%, from 3.1 *±* 0.6 Hz in control to 3.7 *±* 0.6 Hz (n=9, figure 5). OXT had no significant effect on CVNs in the NA (not shown).

**Figure 5.**
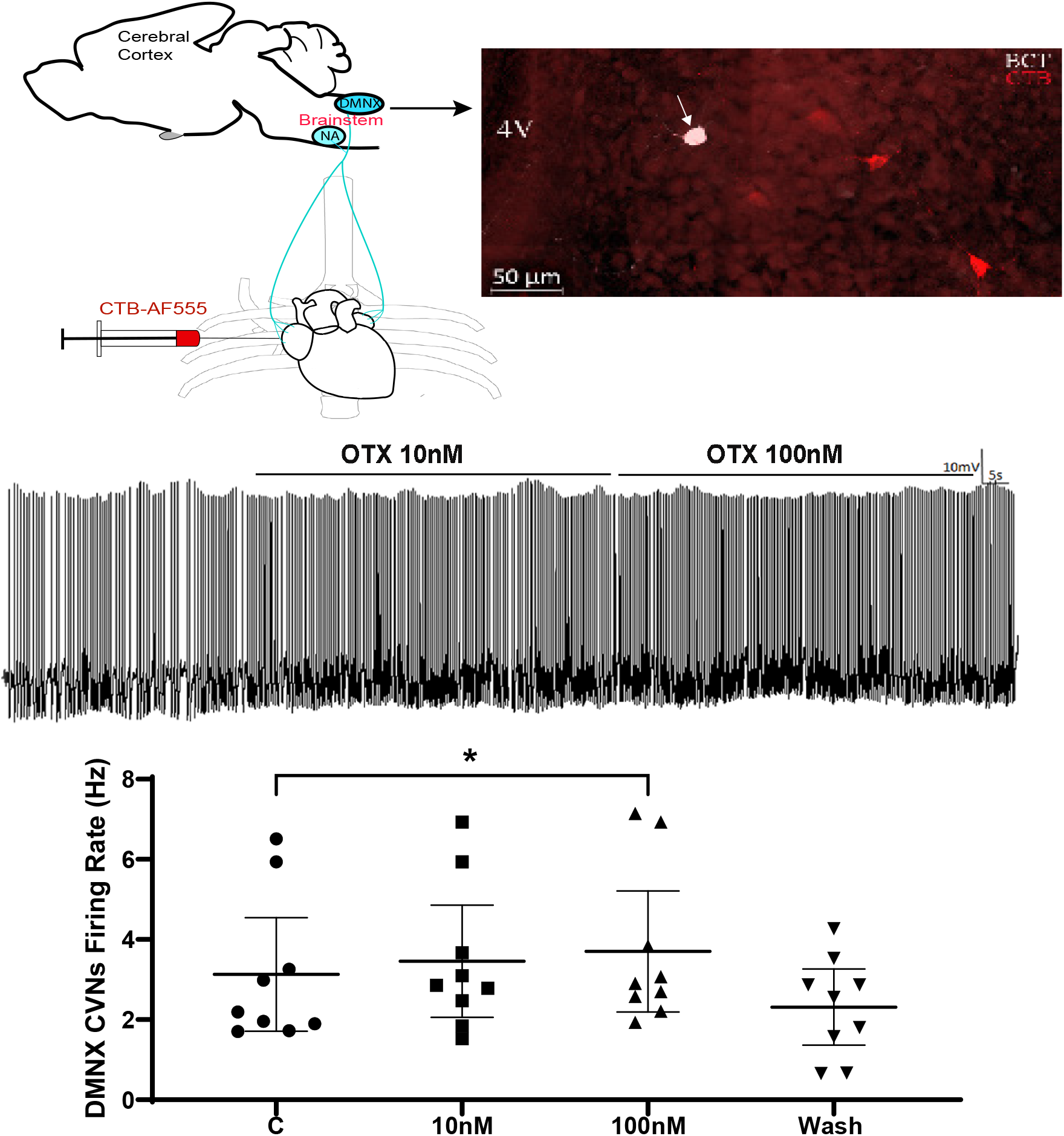
Administration of OXT enhanced CVNs action potential in the brainstem slice. Top left diagram illustrating procedure of retrograde labeling CVNs in the brainstem, top right an example confocal image presenting CTB-555 labeled CVNS (red) and patched CVN neuron (red cell filled with biocytin in white); Middle typical spike trace showing OXT increased CVN neuron action potential firing rate; Bottom bar graph presenting original data with 95% CI (n=9). *p*<*0.05.

## 3.3 Chemogenetic activation of DMNX OXTR+ CVNs increases parasympathetic activity to the heart

To dermine whether excitation of OXTR+ CVNs in the DMNX could alter HR, DREADDs were expressed in OXTR+ CVNs using a retrograde Cre dependent site-specific recombinase FlpO virus (addgene AAV 87306) injected into the pericardial sac, followed 4 weeks later with an injection of a Flp-dependent DREADDs virus (Addgene AAV 154868 -pAAV-hSyn-fDIO-hM3D(Gq)-mCherry) into the DMNX. IP injection of the DREADDs agonist CNO (1 mg/kg) evoked a rapid and long-lasting bradycardia (figure 6), with a maximum HR reduction of 53 *±* 8 bpm observed one hour after CNO injection (p*<*0.01). There were no significant changes in RR in any group. There were no significant changes in HR with CNO in non-DREADDs expressing animals or with saline injections in DREADDs expressing animals.

**Figure 6.**
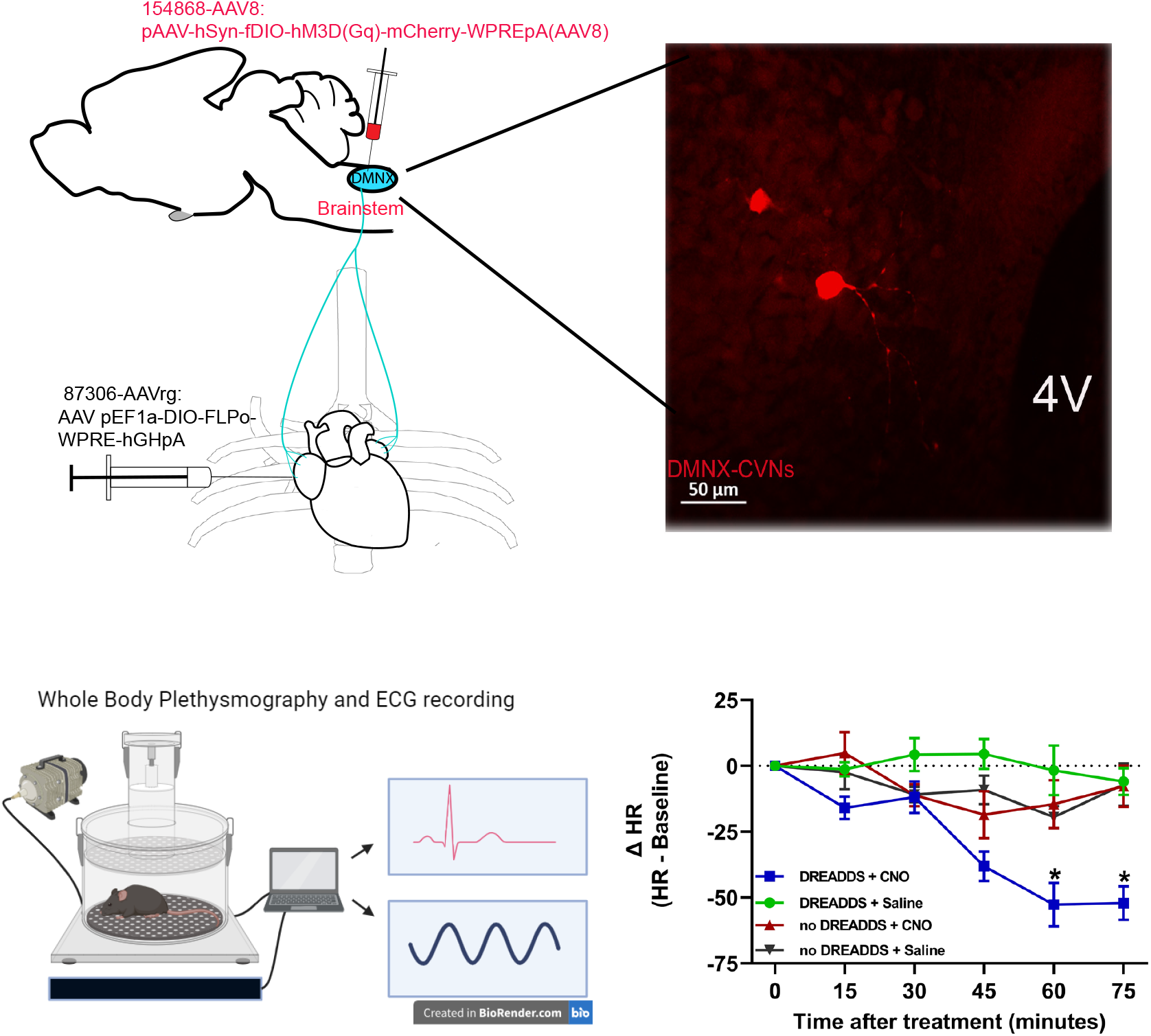
Chemogenetic activation of OXTR+ CVNs increases parasympathetic activity to the heart. Upper left diagram illustrating procedure of two-step viral vector injection to specifically identify and activate CVNs that project to the heart; top right an example confocal image showing cells with mCherry expression in the brainstem slice (red); bottom left diagram illustrating plethysmography and ECG recording and bottom right presenting the statistic results. *p*<*0.05.

## 4 DISCUSSION

There are four major findings from this study. Over half of the CVNs in the DMNX co-localize with OXT receptor positive neurons. Surprisingly, CVNs in the NA, as well as other ChAT neurons in the NA, have sparse expression of OXT receptors. OXT increased the firing of CVNs in the DMNX, but had no effect in CVNs in the NA. Selective chemogenetic activation of OXTR+ CVNs in the DMNX elicited a rapid and long-lasting decrease in HR.

A long-standing issue in the field is whether the different populations of CVNs in the NA and DMNX play overlapping or different physiological roles. Anatomical work has shown CVNs originating from both the NA and DMNX project to cardiac ganglia where their axons form extensive basket endings around ganglionic principal neurons (Cheng et al. (1999)). However, more recent studies found that neurons in the NA and DMNX innervate the same cardiac ganglia but target different subpopulations of principal neurons within the ganglia (Cheng et al. (2004)). More specifically, fibers from the DMNX were found to project to small intensely fluorescent cells (possibly interneurons) as well as to cardiac principal neurons, whereas axons from NA neurons innervate only principal neurons. Lesions of the NA almost completely abolished the baroreflex control of the HR, whereas lesions of the DMNX did not, suggesting that the NA plays a more important role in baroreflex control of HR than the DMNX (Cheng et al. (1999)). Other work found the majority of DMNX CVNs have no obvious input from central or peripheral respiratory or cardiac-related inputs but are activated by stimulation of pulmonary C fiber afferent fibers, suggesting DMNX CVNs may play a more important role in pulmonary C-fiber-evoked bradycardia (Jones et al. (1998)). It is also worth noting that, using chemogenetic activation, long-term stimulation of only CVNs in the DMNX reduced blood pressure in the spontaneously hypertensive rat (Moreira et al. (2018)).

OXT is only synthesized in a limited number of discrete brain regions: the PVN, and supraoptic and accessory nuclei of the hypothalamus (Son et al. (2022)). While many brain areas receive dense projections from OXT neurons, and a multitude of neuronal populations strongly express OXT receptors, there is surprisingly little spatial correlation between OXT fibers and receptors. Only a few neuronal populations both receive dense OXT fibers and have robust expression of OXT receptors. The medulla has the highest positive correlation ratio of OXT fibers to OXT receptors in the CNS (Son et al. (2022)).

Prior work from our lab, and others, have shown there are dense projections from PVN OXT neurons to nuclei in the medulla, and more specifically, there is a monosynaptic excitatory pathway from PVN OXT neurons to CVNs in the DMNX, and that activation of the fibers in this pathway releases OXT (Pinol et al. (2012), Piñol et al. (2014), Jameson et al. (2016), Garrott et al. (2017), Dyavanapalli et al. (2020b), Rodriguez et al. (2023), Schunke et al. (2023)). Neurons in the DMNX have previously been shown to have a robust expression of OXT receptors (Son et al. (2022)). The results in this study advance this work by demonstrating a majority of CVNs in the DMNX are among only a few neuronal populations that receive dense OXT fibers that release OXT and have robust expression of post-synaptic OXT receptors. OXT increases the firing of OXTR+ CVNs in the DMNX, and chemogenetic activation of OXTR+ CVNs decreases HR. Future work is needed to test if selective activation of OXTR+ CVNs in the DMNX prevents or reverses autonomic imbalance and dysfunction in cardiovascular and other diseases. Additionally, as a unique population of OXTR+ CVNs in the DMNX has been identified that when stimulated decreases HR, additional work is needed to identify co-activators of this specific sub-population of OXTR+ CVNs.

## ACKNOWLEDGMENTS

This work was supported by NIH grant R01HL147279-01, NIH Shared Instrument Grant program NIH 1S10OD028515-01, NIGMS P20 GM113134 and Hawaii Community Foundation grant MedRes 2023 00002772.

